## Supplemental Material for "*wnt16* regulates spine and muscle morphogenesis through parallel signals from notochord and dermomyotome"

### SUPPLEMENTAL FIGURES

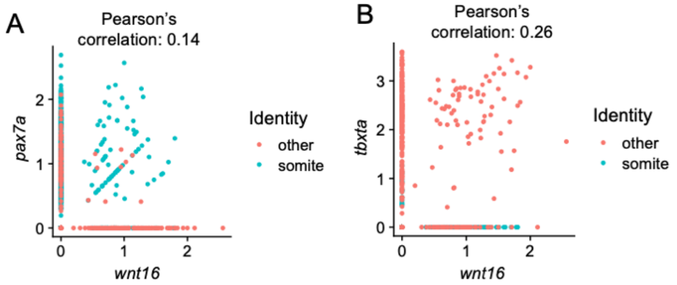


Supplemental Fig 1. *wnt16* expression is differentially co-expressed with *pax7a* and *tbxta* in cells within the “somite” and “other” clusters, respectively.

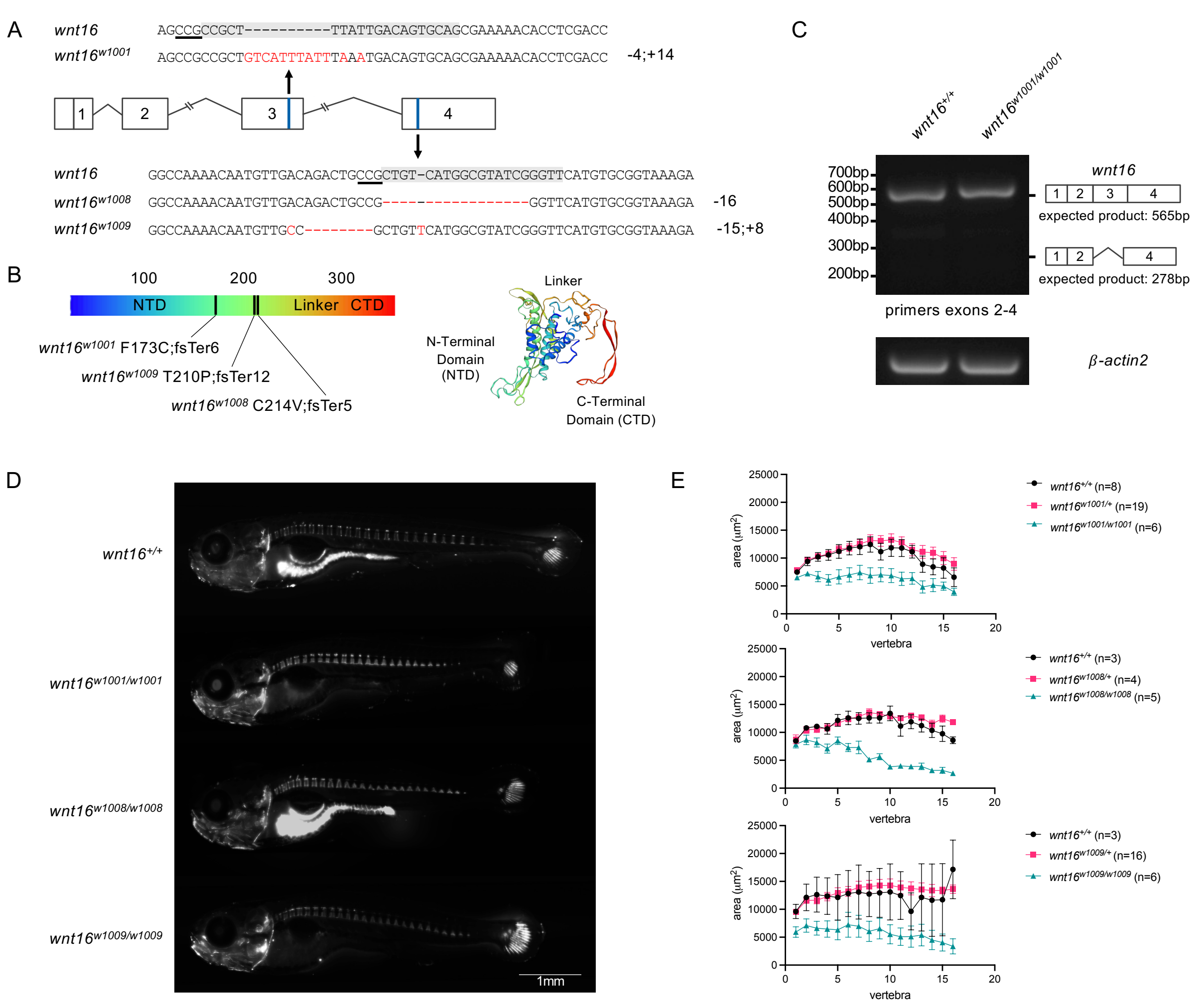


Supplemental Fig 2. Isolation of *wnt16* mutant alleles. (A) Sequence and genomic location of *w1001*, *w1008*, and *w1009*. Grey highlight indicates gRNA target sequence used for CRISPR-based gene editing with PAM underlined. (B) Predicted effects of alleles on amino acid sequence. (C) RT-PCR assessing *wnt16* transcript in *wnt16*$^{w1001}$ mutants. No evidence of transcript reduction or alternative splicing is observed. (D) Calcein staining 13 dpf animals show similar reductions in vertebral mineralization and post-cranial body length in *w1001*, *w1008*, and *w1009* mutants. (E) Quantification of mineralized area shows similar changes in mutants for all three alleles.

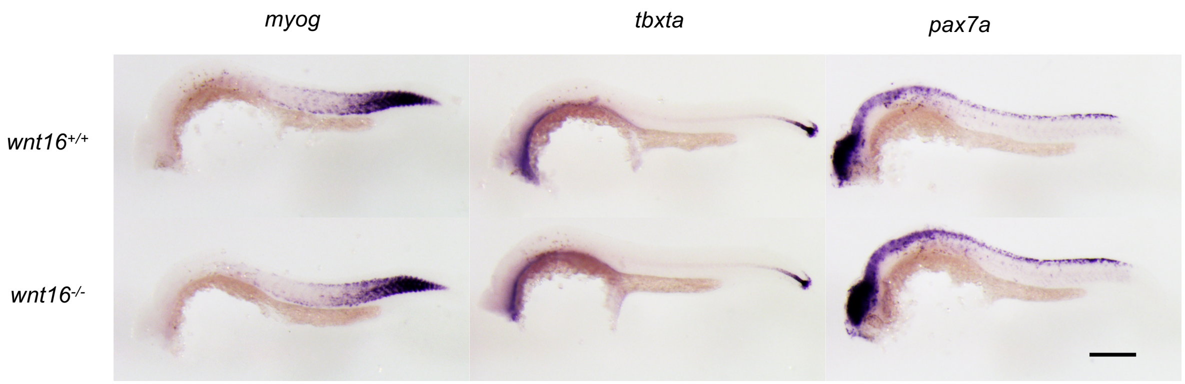


Supplemental Fig 3. RNA ISH for markers of early muscle (*myog, pax7a*) and notochord (*ntla/tbxta*) differentiation in *wnt16*$^{-/-}$ mutant embryos at 1 dpf. Scale bar: 200 μm.

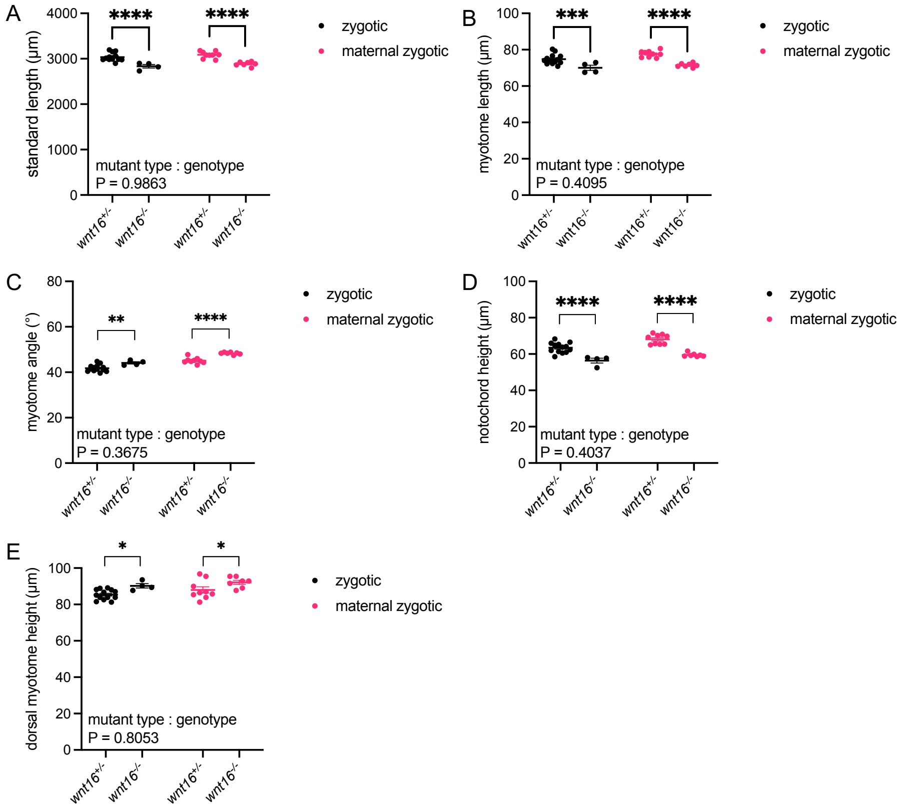


Supplemental Fig 4. Zygotic and maternal zygotic *wnt16*$^{-/-}$ mutant phenotypes. P-values were determined using a two-way ANOVA with Fisher's LSD post hoc test. *p<0.05, **p<0.01, ***p<0.001, ****p<0.0001.

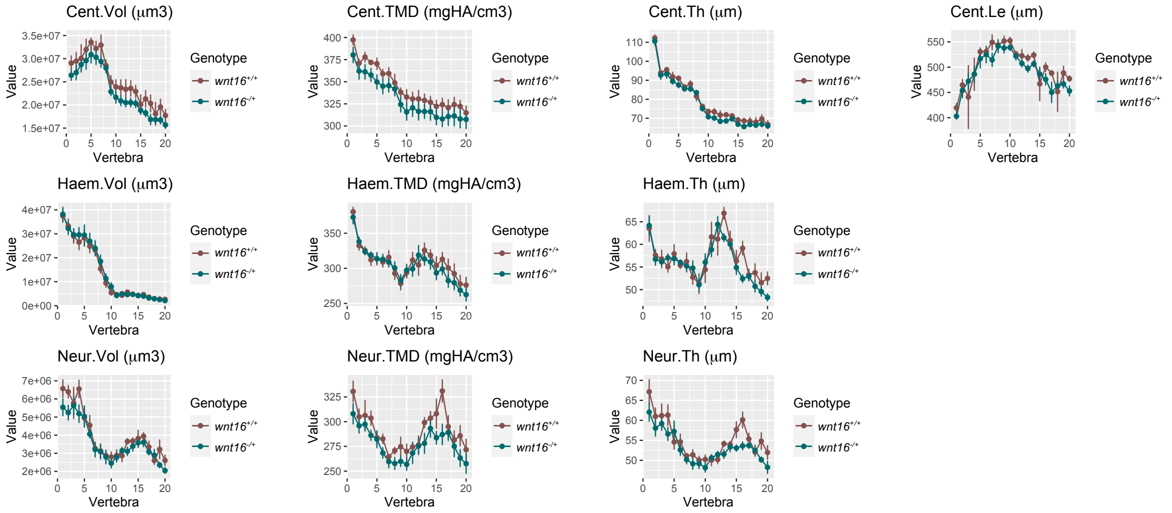


Supplemental Fig 5. Heterozygous *wnt16*$^{-/+}$ mutants do not exhibit significant differences in bone measures compared to wildtype clutchmates.

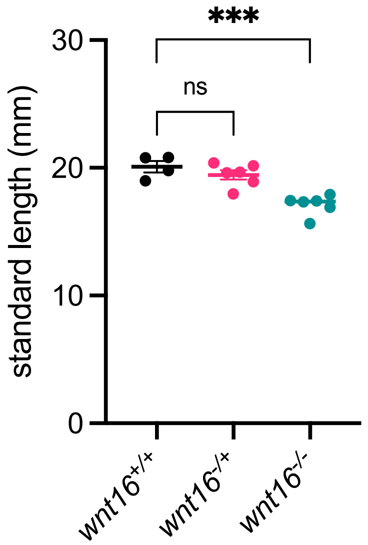


Supplemental Fig 6: *wnt16*$^{-/-}$ mutants exhibit reduced standard length compared to wildtype and heterozygous clutchmates. P-values were determined using one-way ANOVA with Fisher's LSD post hoc test. ***p<0.001, ns: not significant.

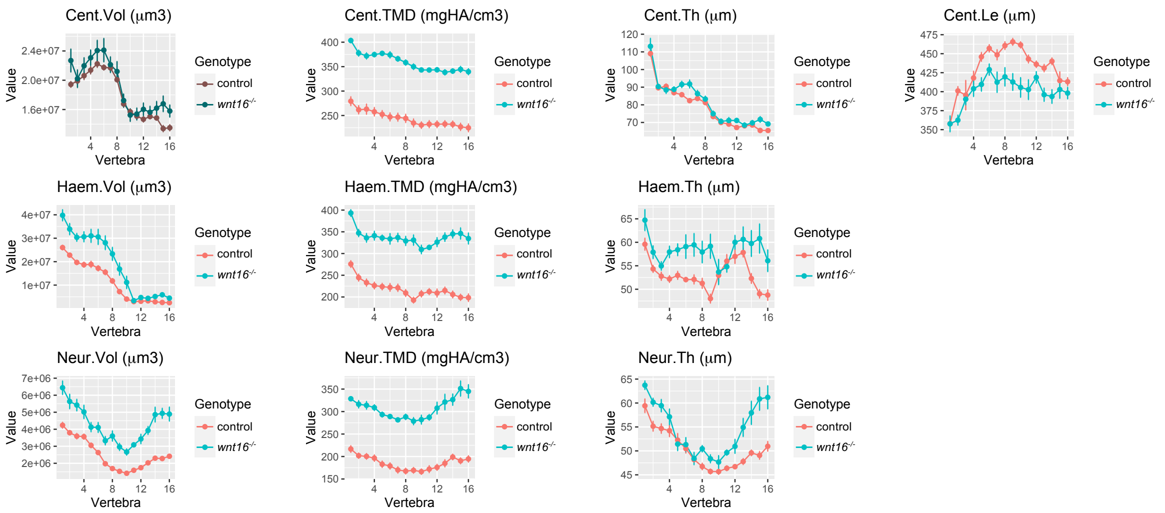


Supplemental Fig 7: Analysis of *wnt16*$^{-/-}$ fish following allometric normalization for standard length. *wnt16*$^{w1001/w1001}$ mutants exhibit significant differences for most measures (all except for Cent.Vol) when normalized for differences in standard length.

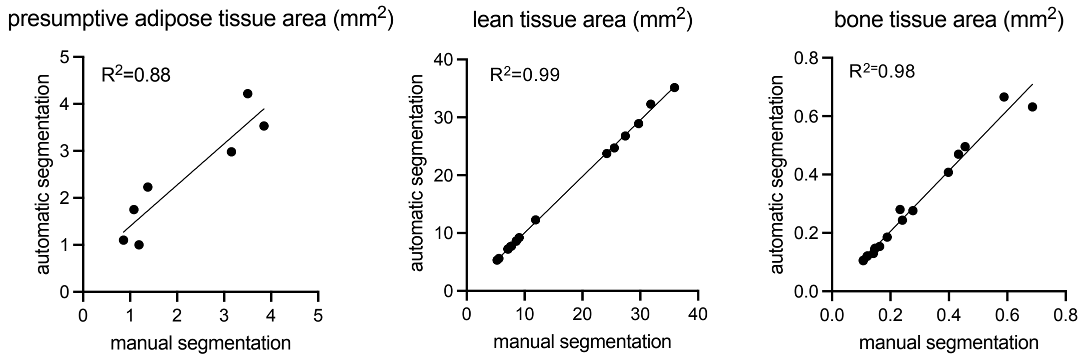


Supplemental Fig 8. Correlation in tissue area computed using automatic and manual segmentation for presumptive adipose tissue (left), lean tissue (middle), and bone tissue (right).

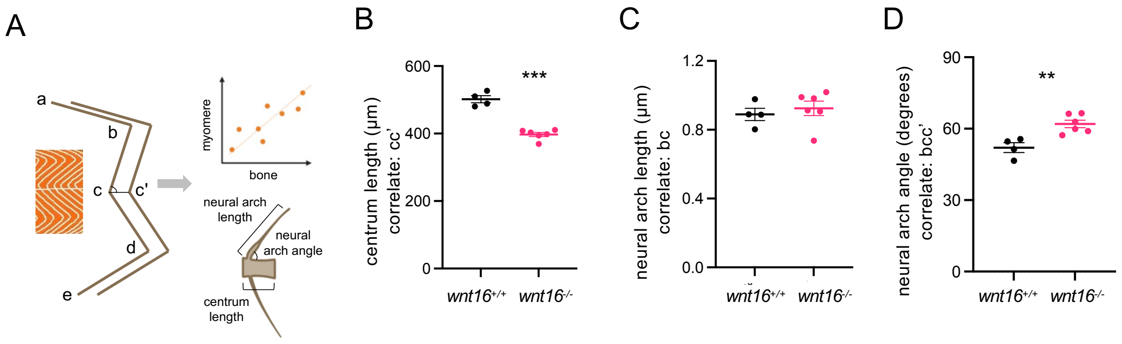


Supplemental Fig 9. Inference of myomere morphology from vertebral measures. (A) Schematic demonstrating correlated features. (B-D) *wnt16*$^{-/-}$ mutants exhibit altered centrum length and neural arch angle. P-values were determined using an unpaired t-test. $^{**}p<0.01$, $^{***}p<0.001$.

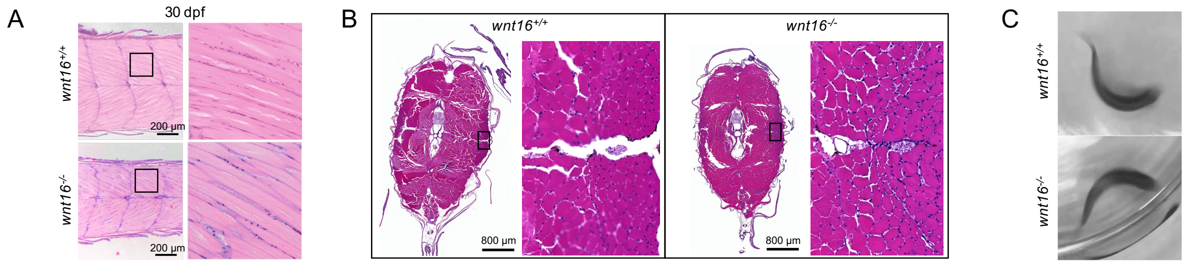


Supplemental Fig 10. *wnt16*$^{-/-}$ mutants do not exhibit obvious muscle pathology. H&E-stained sections showing frontal section of the caudal region and corresponding magnified areas in 30 dpf animals (A) and transverse sections in adult animals (B). *wnt16*$^{-/-}$ mutants exhibited no obvious differences in muscle segmentation or fiber morphology. (C) Mutants and wildtype clutchmates exhibited full-length body flexions when the startle-induced C-start response was evoked.

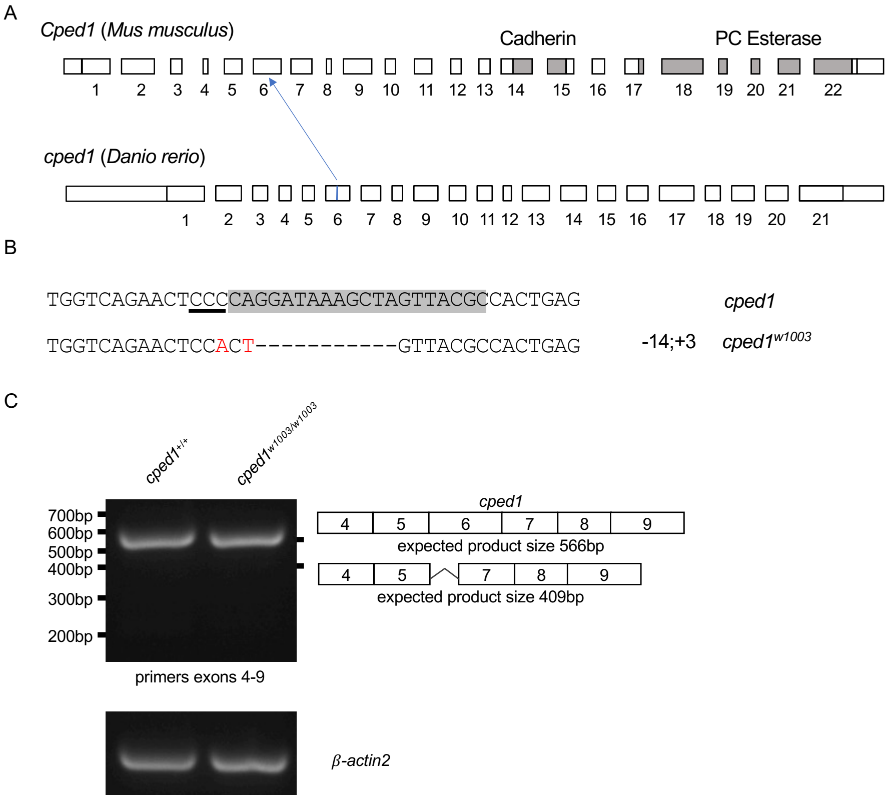


Supplemental Fig 11. Isolation of *cped*$^{w1003}$. (A) Genomic location of *w1003* and its corresponding location mapped to mouse *Cped1*. Locations of Cadherin and PC Esterase domains in mouse *Cped1* are from (Maynard et al., 2018). (B) Genomic sequence of *w1003*. Grey highlight indicates gRNA target sequence used for CRISPR-based gene editing with PAM underlined. (C) RT-PCR assessing *cped1* transcript in *cped1*$^{w1003}$ mutants. No evidence of transcript reduction or alternative splicing is observed.

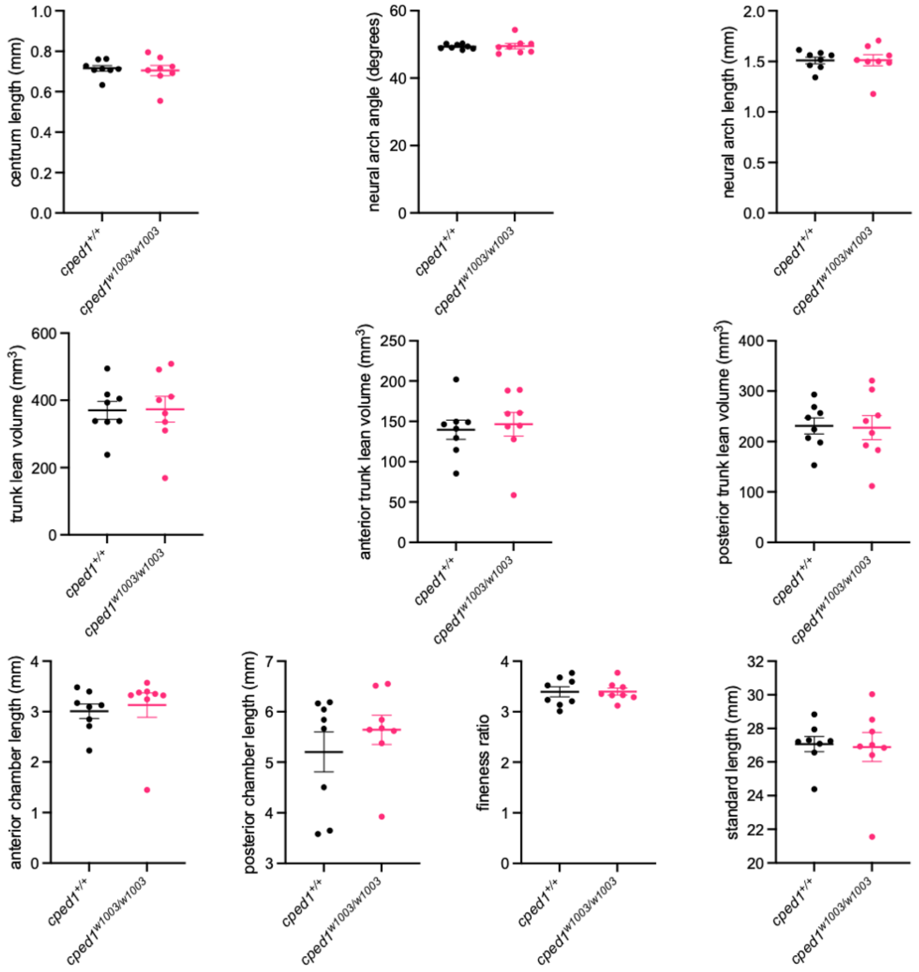


Supplemental Fig 12. Data used for calculating Z-scores for Fig 9D.

### Sample MATLAB code used for computing lean tissue volume

```
v = dicomreadVolume('mutant 01');

v = squeeze(v);
thresh1 = 0;
thresh2 = 1043;
thresh3 = 6368;
startk = 236;
endk = 822;

dims = size(v);
imax = dims(1);
jmax = dims(2);
kmax = dims(3);

% Target A = noise less than 0
targetA = v;

% Target B = noise between 0 and soft tissue
targetB = v;

% Target C = soft tissue ie muscle
targetC = v;

% Target D = bone
targetD = v;

for i = 1:imax
    for j = 1:jmax
        for k = 1:kmax
            p = v(i, j, k);
%target A
           if(p < thresh1)
                targetA(i, j, k) = 1;

            else
               targetA(i, j, k) = 0;
           end
%target B
           if((p > thresh1) && (p < thresh2))
                targetB(i, j, k) = 1;

            else
               targetB(i, j, k) = 0;
           end
%target C
           if((p > thresh2) && (p < thresh3))
                targetC(i, j, k) = 1;

            else
               targetC(i, j, k) = 0;
           end
%target D
           if(p > thresh3)
                targetD(i, j, k) = 1;

            else
               targetD(i, j, k) = 0;
           end
        end
    end
end

%cross section
csA(k) = 0;
csB(k) = 0;
csC(k) = 0;
csD(k) = 0;
```

```
% output is in mm^2
for k = 1:kmax
    csA(k) = sum(targetA(:,:,k), 'all') * 0.000441;
    csB(k) = sum(targetB(:,:,k), 'all') * 0.000441;
    csC(k) = sum(targetC(:,:,k), 'all') * 0.000441;
    csD(k) = sum(targetD(:,:,k), 'all') * 0.000441;
end

% output is in mm^3
volA = (sum(csA) * 0.021);
volB = (sum(csB) * 0.021);
volC = (sum(csC) * 0.021);
volD = (sum(csD) * 0.021);

headlesscsA = csA(startk:endk);
headlesscsB = csB(startk:endk);
headlesscsC = csC(startk:endk);
headlesscsD = csD(startk:endk);

% output is in mm^3
headlessvolC = (sum(headlesscsC) * 0.021);
headlessvolD = (sum(headlesscsD) * 0.021);
```
